# The route of infection shapes Rift Valley fever virus pathogenesis, humoral immune response, and horizontal transmission in sheep

**DOI:** 10.64898/2026.03.12.711297

**Authors:** Sara Morán de Bustos, Iris Sánchez-Del Pozo, Miriam Pedrera, José J. Cerón, Elisabeth Fuentes, David Sardón, David Rodríguez-Temporal, Belén Borrego, Alejandro Brun, Belén Rodríguez-Sánchez, Pedro J. Sánchez-Cordón

## Abstract

Rift Valley fever (RVF) is a zoonotic arboviral disease that causes adverse pregnancy outcomes and high mortality in domestic and wild ruminants. The disease is caused by the RVF virus (RVFV), which is transmitted by mosquitoes from several genera, mainly *Aedes* and *Culex*. However, whether ruminants can become infected by horizontal virus transmission remains unclear. In addition, how the route of RVFV inoculation may influence RVF pathogenesis and the host immune response in animals is still largely unknown. With this aim, we conducted a comparative experimental study in which young sheep were either inoculated subcutaneously (SC) or intranasally (IN) with the virulent RVFV 56/74 strain. We then evaluated disease dynamics, viremia, virus excretion, tissue damage, and the humoral immune response. We also aimed to determine whether RVFV can be transmitted from infected to in-contact animals, and to assess whether the inoculation route may influence virus excretion and the likelihood of subsequent horizontal transmission. The results showed that SC inoculated sheep had a shorter incubation period, an earlier onset of viremia, and an earlier seroconversion. In contrast, IN inoculated animals developed higher rectal temperatures, reached higher peak viremia, and developed a more robust neutralizing antibody response. They also exhibited increased concentrations of analytes indicative of moderate but more severe hepatic injury compared with the subcutaneous group, along with more pronounced histopathological damage in the central nervous system. These results demonstrate the influence of the route of inoculation on RVF pathogenesis and the host immune response. Our results also confirmed the horizontal transmission of RVFV between SC inoculated sheep and in-contact animals housed in the same room, a phenomenon not observed in the IN inoculated group. This finding underscores the influence of the inoculation route on virus transmission and the potentially significant role of horizontal transmission in RVF epidemiology and disease control.

**Author summary:** According to the World Health Organization (WHO), RVFV is considered a priority pathogen due to its ability to strain animal and public health systems, especially in developing countries. RVF outbreaks have occurred across most of Africa and, since 2000, in the Arabian Peninsula. Evidence of RVFV circulation in North Africa further highlights the threat to Europe, where competent mosquito vectors are present. How the inoculation route shapes disease dynamics and hosts immunity is still largely unknown. Similarly, whether the virus can spread between infected and non-infected animals without competent vectors remains unclear. A comparative infection in which young sheep were inoculated SC or IN with the RVFV 56/74 strain showed that SC inoculated sheep had a shorter incubation period, an earlier onset of viremia, and earlier seroconversion. However, rectal temperature and peak viremia were higher in IN inoculated sheep, which also showed evidence of moderate but more severe hepatic damage, accompanied by greater central nervous system damage. Only the in-contact animals housed in the subcutaneous group became infected, demonstrating horizontal transmission. Our results show that the route of inoculation influences disease progression and that RVFV can be transmitted among sheep in the absence of mosquitoes.

## Introduction

Rift Valley fever (RVF) is a zoonotic arboviral disease that causes adverse pregnancy outcomes and high mortality in livestock, including cattle, sheep, camels and goats as well as in wild ruminants [1,2]. The disease is caused by the Rift Valley fever virus (RVFV), an RNA virus belonging to the family *Phenuiviridae* (genus *Phlebovirus*), which is transmitted by mosquitos of different genera, mainly *Aedes* and *Culex*. Despite the existence of multiple lineages, there is a limited genomic variability among RVFV strains, with only one serotype recognised so far [3]. In humans, direct transmission of RVFV is considered the primary route of infection and may occur through exposure of wounds or mucosal surfaces with infectious animal tissues, body fluids, or aborted material [4,5]. Serological data suggests that the virus circulates in nearly all African countries [6]. Since 2000, RVF outbreaks have been reported in the Arabian Peninsula and on islands off the coast of Southern Africa, with more recent events documented in Kenya, Mayotte Island, Sudan, and Uganda [7]. Further outbreaks were reported in 2025 in Senegal and Mauritania, where, in addition to the devastating consequences for livestock production, 42 human deaths have been recorded [8]. Countries in the Mediterranean basin, such as Turkey, Tunisia and Libya, have also reported serological evidence of RVFV circulation [9,10], highlighting the potential for the virus to spread and the threat it poses to Europe, where mosquito species competent for virus transmission are already present [11]. In this sense, the RVFV has been catalogued a priority pathogen requiring urgent R&D investment [12] due to its potentially devastating consequences for livestock production and public health systems, particularly in developing countries [13].

Host susceptibility to RVFV varies by species and age, with sheep and goats generally more affected than cattle or camels, and lambs being particularly vulnerable, showing mortality rates that can reach 90–100%. In contrast, adult sheep tend to show milder disease, typically developing pyrexia and nonspecific clinical signs, so abortion storms, stillbirths and foetal malformations are often the only indicators of infection in pregnant animals [14,15]. RVFV can also cause infection and outbreaks in humans, with symptoms ranging from mild disease to a severe haemorrhagic form that is often fatal [16]. Association between RVFV infection and miscarriage in pregnant women has also been suggested, although the underlying mechanisms remain poorly understood [17]. No vaccines against RVFV are available for human use and, though licensed veterinary vaccines are available, these have major drawbacks, including abortion that limit their use [18]. As a result, considerable efforts are being directed toward the development of new, safer, and more effective vaccines [18–20]. However, studies aimed at understanding the pathogenic mechanisms of RVFV and the host immune responses in susceptible livestock species remain scarce, despite this knowledge being a critical prerequisite for developing and improving next-generation vaccines. Experimentally, in addition to factors such as species and age, other variables like the inoculation route should also be considered [21]. In this regard, it remains largely unknown how the route of RVFV infection may influence pathogenesis, the activation of the host immune response, and ultimately the disease outcome in livestock. For this purpose, sheep constitute an excellent experimental model, playing a key role not only as reservoir hosts but also in the transmission of RVFV to humans, which allows findings from RVFV studies in sheep to be extrapolated to other ruminant species. So far, the RVFV infection model in sheep has not been studied in detail, leaving key questions unresolved. These include how the route of infection may influence disease dynamics, viremia, viral dissemination across organs and virus excretion, viral replication and virus–cell interactions in target tissues, the extent of tissue damage, and the resulting humoral and cellular immune responses.

Infection models for pathogenesis studies in sheep have usually used the subcutaneous, intravenous or intraperitoneal route [21]. However, unlike some studies conducted in goats and cattle [22,23], intranasal infections have not been performed in sheep, even though RVFV could also be transmitted through aerosol exposure[24,25]. Moreover, to date, no comparative experimental infections using intranasal and parenteral routes (intramuscular, intradermal, or subcutaneous) have been conducted in sheep. Some experimental studies have also reported sporadic cases of horizontal transmission to contact-exposed sheep [26–28], whereas others failed to demonstrate horizontal transmission from intravenously infected lambs to either naïve or immunosuppressed lambs [29]. Therefore, it remains unclear whether horizontal transmission constitutes a natural mode of infection among animals within herds during RVF outbreaks, or what role it plays in RVFV epidemiology.

In the present study, we aimed to address a dual objective. On the one hand, we sought to determine how the route of RVFV inoculation (intranasal vs. subcutaneous) may influence disease dynamics, viremia, virus excretion, tissue damage, and the humoral immune response in sheep. On the other hand, we aimed to determine whether RVFV transmission occurs between infected and in-contact animals, and whether the inoculation route may influence virus excretion and subsequent transmission.

## Results

### Disease dynamics and macroscopic evaluations

Seven young sheep per experimental group were inoculated subcutaneously (SC) or intranasally (IN) with RVFV 56/74 strain. Additionally, three animals within each group were mock inoculated and designated as in-contact animals. Throughout the experiment (Fig. 1), none of the animals inoculated either SC or IN showed overt clinical signs. Despite the subclinical presentation of the disease, all animals in the SC inoculated group, except sheep #11 and #17, showed an earlier increase in rectal temperature between 1 and 2 days post-inoculation (dpi), reaching peaks of up to 41.6 °C (Fig. 2A). In contrast, all sheep IN inoculated showed an increase in rectal temperature, with peaks up to 41.9 °C between 2 and 3 dpi (Fig. 2B). Statistical analysis revealed that, at 2 and 3 dpi, temperatures were significantly higher in the IN inoculated sheep than in the SC inoculated group (Fig. 2C). Thus, while the incubation period was shorter in the SC inoculated animals, the IN inoculated sheep exhibited higher temperature peaks.

**Fig. 1.**
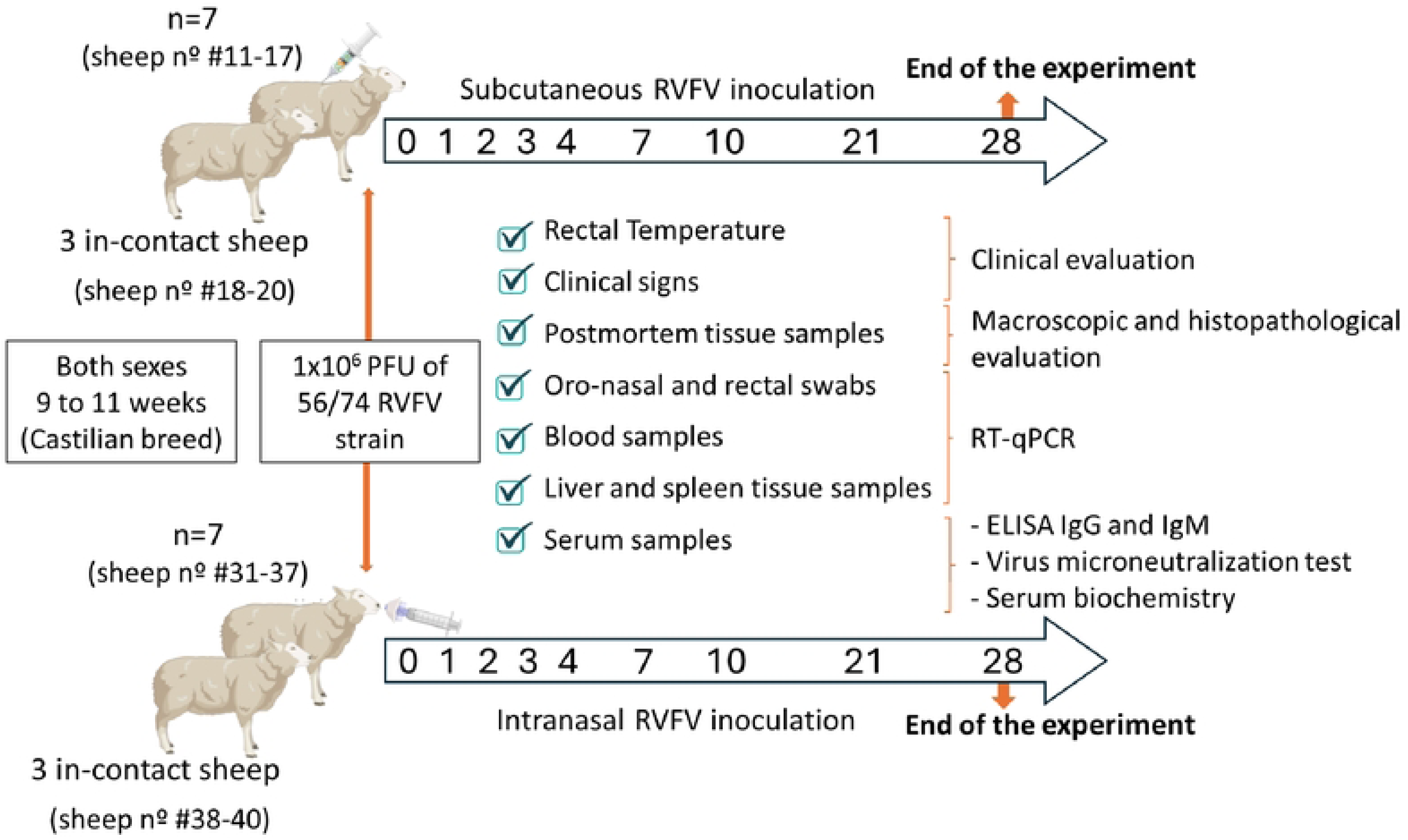
Animal experimental design. Twenty sheep of both sexes, 9 to 10 weeks old were randomly assigned into two groups with ten animals each. In each group, seven animals were inoculated subcutaneously or intranasally with 1 mL containing 1 × 10^6^ PFU/mL of the virulent RVFV 56/74 strain. Additionally, three animals within each group were inoculated with mock-infected cell culture medium and designated as “in-contact animals”. Following inoculation, rectal temperature and clinical signs were monitored daily. Oro-nasal and rectal swabs, as well as EDTA blood and serum samples were collected from all the animals prior to infection (day 0) and at 1, 2, 3, 4, 7, 10, 21 and 28 days post-infection (dpi). At the end of the experiment (28 dpi), all animals were euthanised and necropsied. During necropsies, macroscopic lesions were scored and tissue samples collected. Image created with Biorender.com.

**Fig. 2.**
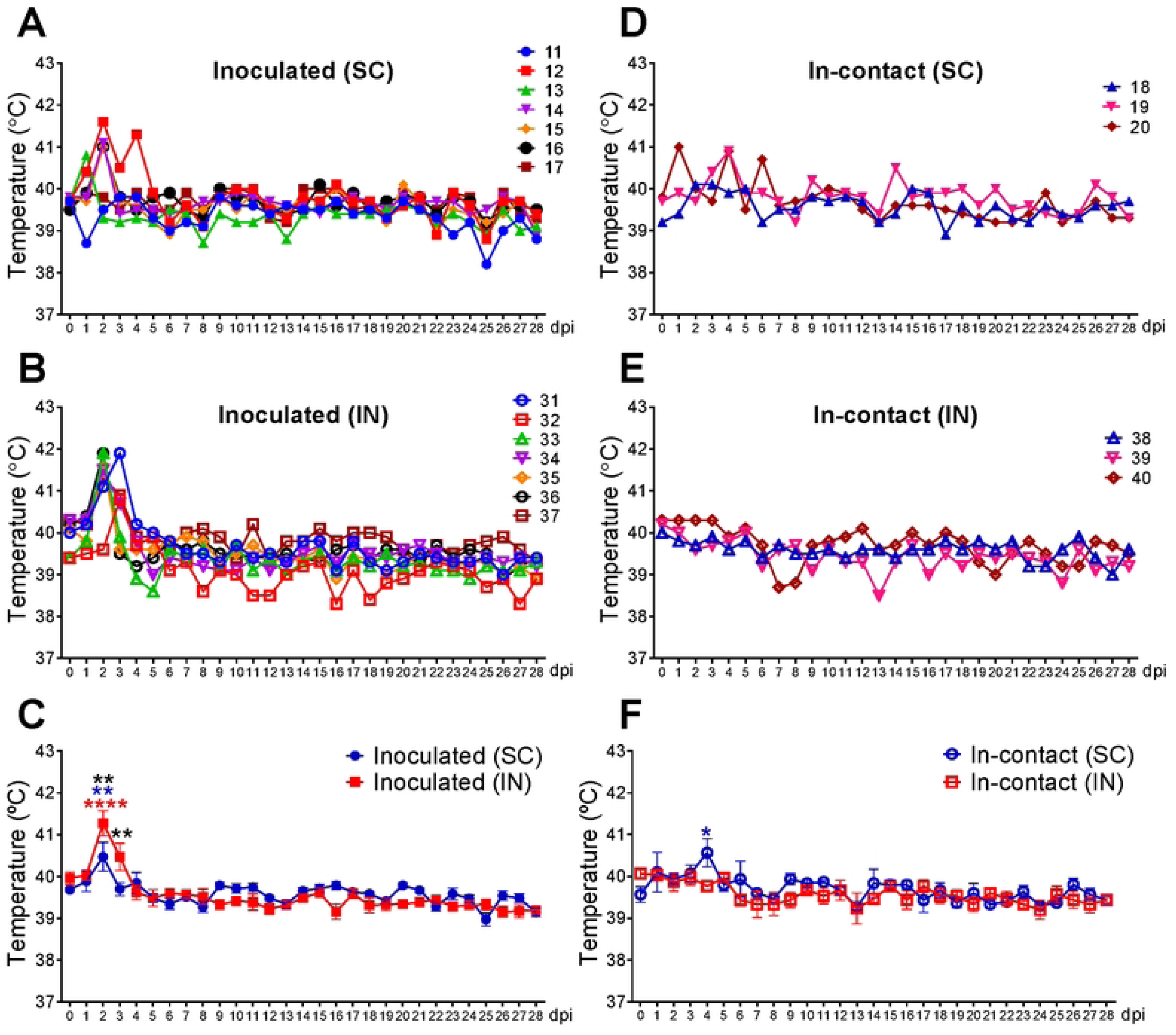
Kinetics of rectal temperature in sheep inoculated with RVFV by different routes and in-contact sheep. Individual kinetics of rectal temperature in sheep inoculated by subcutaneous or intranasal route with RVFV 56/74 strain **(A-B)** and in-contact sheep **(D-E)**. Each symbol represents the values for an individual sheep at the different time points. Statistically significant differences at different days post-inoculation (mean ± SD) between subcutaneously and intranasally inoculated animals **(C)** and between both groups of in-contact sheep **(F)** were analysed by two-way ANOVA. Statistical significance between groups is annotated with black asterisks. Statistically significant differences within each experimental group at different days post inoculation relative to pre-inoculation values were assessed by one-way ANOVA. Statistical differences are indicated by red (intranasal group) and blue (subcutaneous group) asterisks **(C and F)**. (x-axis): day post-inoculation (dpi); (y-axis): temperature; variables of significance (*P ≤ 0.05; **P ≤ 0.01; ***P ≤ 0.001; ****P ≤ 0.0001). SC: subcutaneous group; IN: Intranasal group.

The in-contact animals in both experimental groups also showed no clinical signs. However, the animals in contact with those inoculated by the subcutaneous route showed recurrent, transient increases in rectal temperature between 3 and 6 dpi (Fig. 2D), with peaks statistically significant regarding pre-inoculation values at 4 dpi (Fig. 2F). These changes were not observed in animals in contact with those IN inoculated (Fig. 2E).

Macroscopic evaluation at the end of the experiment (28 dpi) revealed no notable macroscopic lesions in any of the inoculated or in-contact animals.

### Detection of RVFV-specific antibodies and the kinetics of neutralising antibodies

The antiviral nucleoprotein ELISA results revealed that all animals inoculated SC tested positive for IgG and IgM in serum samples, with some of them (sheep #13, #14, #15 and #16) seroconverting from 4 dpi (Fig. 3A). Likewise, all animals inoculated IN tested positive for IgG and IgM, but only from 7 dpi onwards (Fig. 3B). Therefore, the results evidenced earlier seroconversion in animals inoculated by subcutaneous route. Regarding in-contact animals, the three animals within the group inoculated SC tested positive in both tests from 7 dpi onwards (Fig. 3C), while the three in-contact animals in the group inoculated IN tested negative for IgG and IgM (Fig. 3D). These results also revealed that the in-contact animals in the subcutaneous group seroconverted later than the inoculated sheep housed in the same room.

**Fig. 3.**
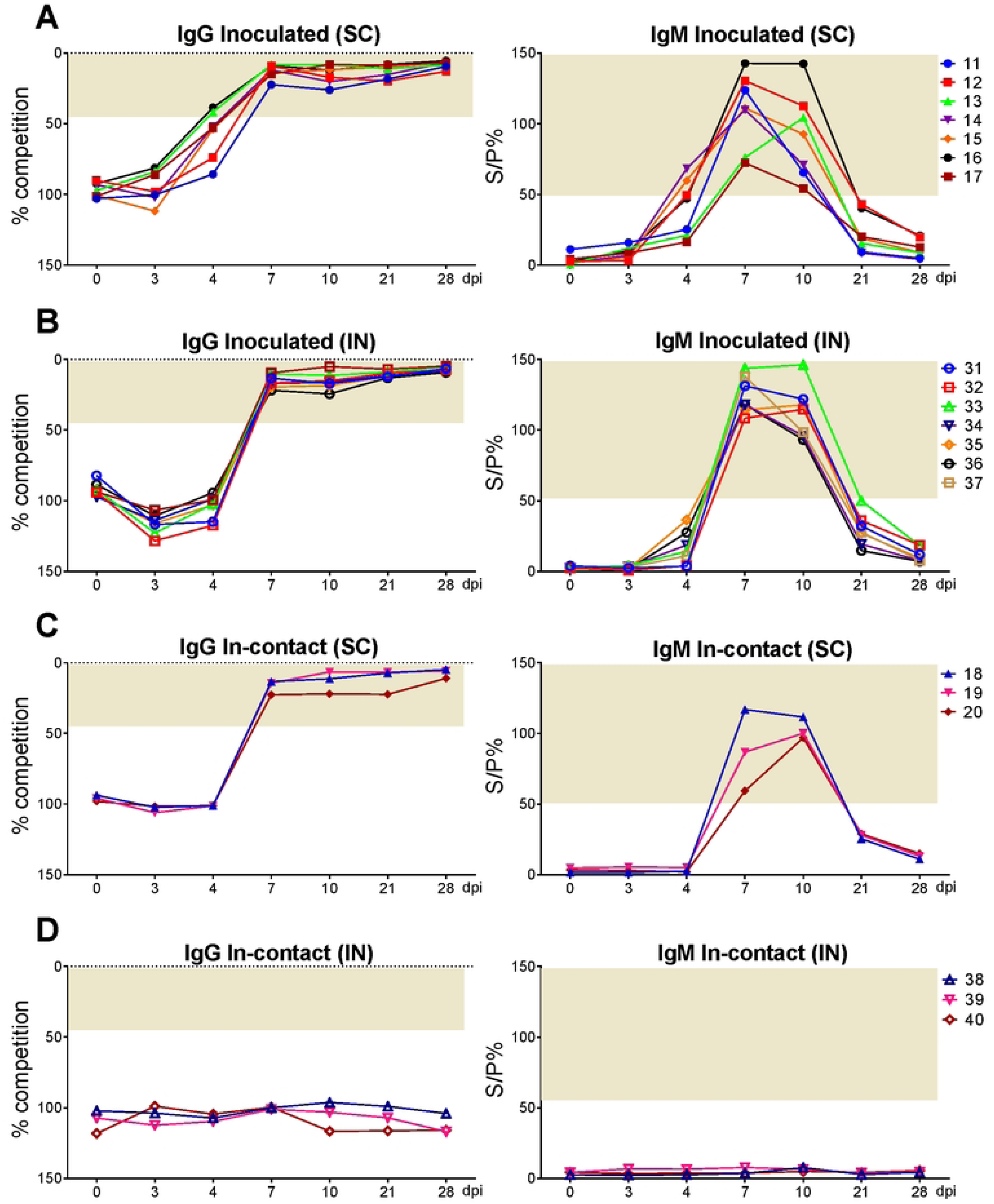
Detection of IgG and IgM against RVFV nucleoprotein. Detection of RVFV-specific IgG and IgM antibodies in serum samples from sheep inoculated by subcutaneous or intranasal route with RVFV 56/74 strain **(A-B)** and in-contact sheep **(C-D)**. Each symbol represents the values for an individual sheep at the different time points. Competition percentage (% competition) and sample-to-positive percentage (S/P%) were determined using the IDVet ELISA test. The grey area indicates the positive threshold defined by the manufacturer. (x-axis): day post-inoculation (dpi); (y-axis): competition percentages and sample-to-positive percentage; SC: subcutaneous group; IN: Intranasal group.

By 4 dpi, all inoculated animals in the subcutaneous group had neutralising antibodies (nAb) titres above the assay threshold values, except for sheep #12, which showed nAb from 7 dpi onwards (Fig. 4A). In the IN inoculated group, all the animals showed a delayed nAb response not detected until 7 dpi. This response was characterised by higher and more homogeneous titres that stabilised earlier into a plateau (Fig. 4B).

**Fig. 4.**
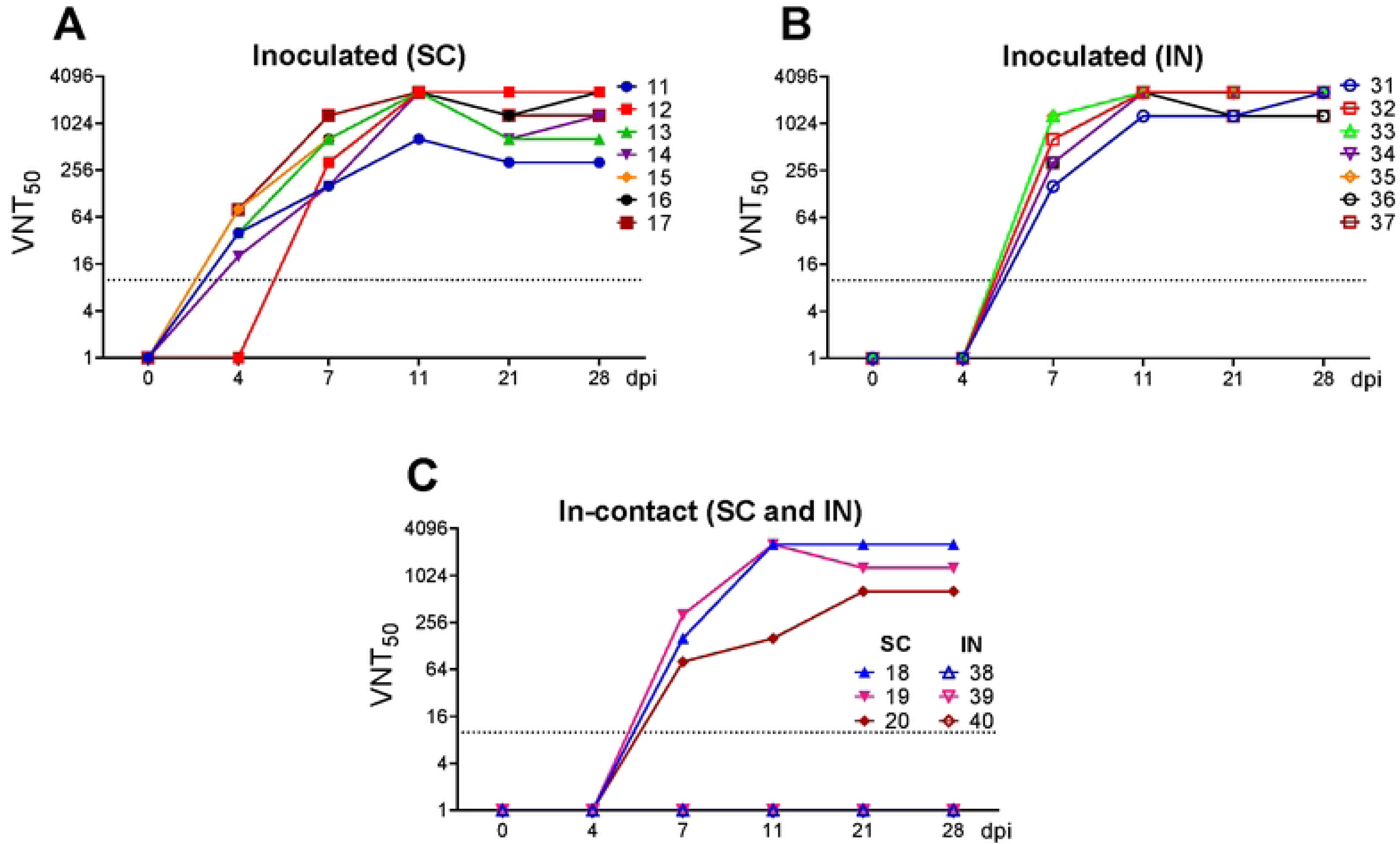
Neutralizing antibody kinetics after RVFV subcutaneous or intranasal inoculation and contact exposure in sheep. Individual kinetics of neutralizing antibody titres in serum samples from sheep inoculated by subcutaneous or intranasal route with RVFV 56/74 strain **(A-B)** and in-contact sheep **(C)**. Each symbol represents the values for an individual sheep at the different time points. The dotted line represents the sensitivity limit of the assay. (x-axis): day post-infection (dpi); (y-axis): reciprocal serum dilution that produces 50% neutralization of cytopathic effect (VNT_50_); SC: subcutaneous group; IN: Intranasal group.

The three in-contact animals in the subcubtaneous group showed nAb by 7 dpi, whereas no nAb were detected at any time in the in-contact sheep in the intranasal group (Fig. 4C). Together with the aforementioned virus specific antibody response, these results substantiate the infection of in-contact sheep in the subcutaneous group, yet not in the intranasal group.

### Viral RNA levels and virus isolation in blood, swabs and tissues

In sheep #17 inoculated SC, the viral genome was not detected in blood samples at any time during the experiment. In the other animals in this group, the viral genome was detected from day 1, with peaks on 1 and 2 dpi (Fig. 5A). In contrast, viral genome was first detected in all IN inoculated sheep from day 2, with peaks on 2 and 3 dpi (Fig. 5B). Regarding in-contact animals in both experimental groups, only those within the group inoculated SC (sheep #18, #19 and #20) showed detectable levels of viral genome from 2 dpi, with peaks between 3 and 4 dpi (Fig. 5D and 5E), evidencing a delay in the onset of viremia compared with the inoculated sheep housed in the same room (Fig. 5C and 5F).

**Fig. 5.**
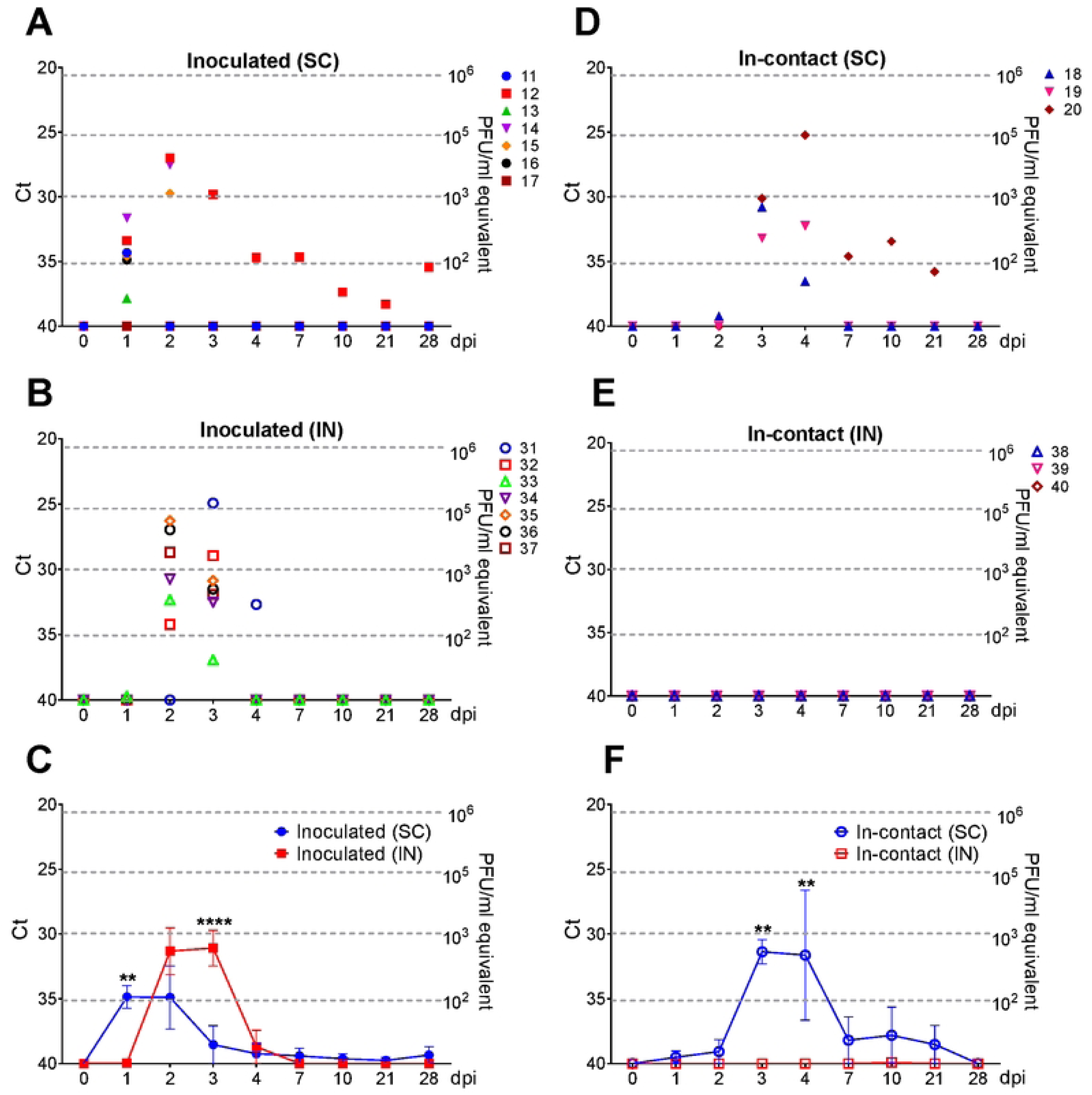
Molecular detection of RVFV 56/74 strain in blood samples. Detection of viral RNA by RT-qPCR in EDTA blood samples from sheep inoculated by subcutaneous or intranasal route with RVFV 56/74 strain **(A-B)** and in-contact sheep **(D-E)**. Each symbol represents the values for an individual sheep at the different time points. Statistically significant differences at different days post-inoculation (mean ± SD) between subcutaneously and intranasally inoculated animals **(C)** and between both groups of in-contact sheep **(F)** were analysed by two-way ANOVA. Statistical significance between groups is annotated with black asterisks. The dotted lines indicate the relation between Ct values and the corresponding infectious viral load (PFU/mL), established using a standard dilution curve generated from samples with known PFU/mL concentrations. (x-axis): day post-inoculation (dpi); (y-axis, left): Ct values; (y-axis, right): plaque-forming units per millilitre (PFU/mL); variables of significance (*P ≤ 0.05; **P ≤ 0.01; ***P ≤ 0.001; ****P ≤ 0.0001); SC: subcutaneous group; IN: Intranasal group.

Attempts were made to isolate the virus *in vitro* from blood samples obtained from inoculated and in-contact animals. For each animal, the sample selected for virus isolation corresponded to the time point showing the highest RNAemia level detected between 1 and 4 dpi. Additional isolation attempts were performed even in animals in which RNAemia was not detected. In sheep that were inoculated, infectious virus was isolated from all selected samples except those from sheep #11, #13 and #17 in the subcutaneous group, and from sheep #33 in the intranasal group, where viral genome was either undetectable or present at very low RNAemia levels (S1 table).

Regarding in-contact sheep housed in the subcutaneous group, virus was isolated from sheep #18 and #20, but not from in-contact sheep #19, which had the lowest RNAemia values. In addition, infectious virus was not isolated from any of the in-contact sheep housed in the intranasal group (S1 table).

On the other hand, virus genome was detected in only four of the seven oro-nasal swabs collected from sheep inoculated IN at 1 dpi, and in one of the seven swabs collected at 2 dpi, although infectious virus could not be isolated from any of these samples. Beyond that, no viral genome was detected in any of the other oro-nasal or rectal swabs taken throughout the experiment, and virus isolation was not attempted.

Eventually, no viral genome was detected in any of the liver or spleen samples collected from either inoculated or in-contact sheep at 28 dpi. Virus isolation was not attempted from these tissue samples.

### Clinical biochemistry

Clinical biochemistry analytes were measured in serum samples to assess the effects of RVFV infection on various organ systems. Concentrations of biochemical markers indicative of liver injury or dysfunction such as serum aspartate aminotransferase (AST) and alkaline phosphatase (ALP) increased moderately in both groups from 2 to 4 dpi. (Fig. 6A and 6B). Individual profiles highlighted the early rise and highest concentrations of these analytes in sheep #12 and #13 (subcutaneous group) and in sheep #31, #33, #34 and #35 (intranasal group). In contrast, gamma-glutamyl transferase (GGT) exhibited a delayed moderate increase beginning at 7 dpi (Fig. 6C), with the IN inoculated animals reaching the highest GGT levels. Therefore, these markers were generally higher in animals inoculated IN, suggesting moderate but greater liver damage or dysfunction compared with the subcutaneous group. Bile acids and bilirubin levels (Fig. 6D and 6E), as well as albumin concentrations (S1 Fig), did not exhibit remarkable changes throughout the experiment.

**Fig. 6.**
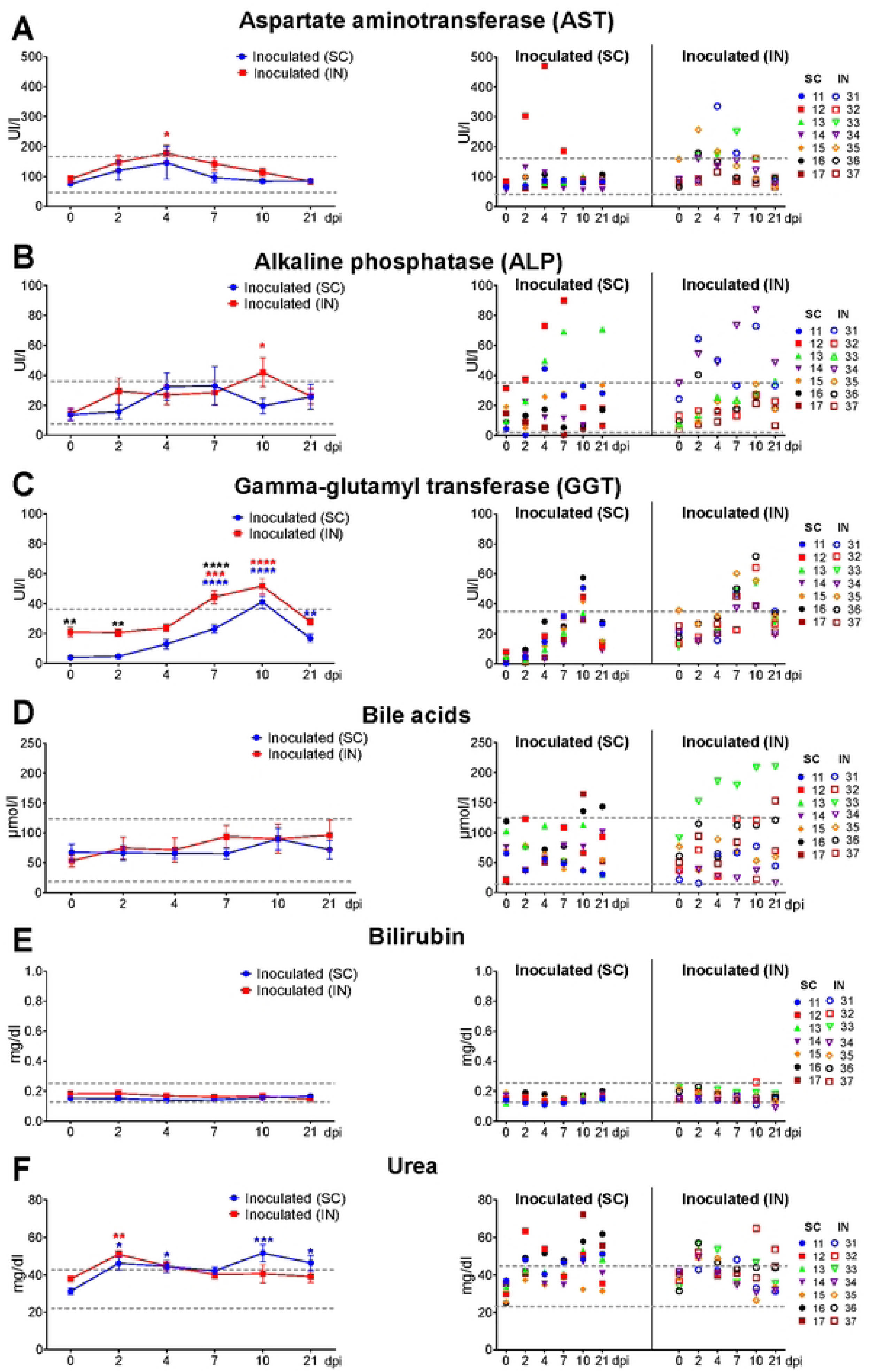
Serum clinical biochemical parameters in sheep inoculated with RVFV by subcutaneous or intranasal route. Serum samples from inoculated sheep were analysed for biomarkers indicative of hepatic damage, including (A) aspartate aminotransferase (AST), (B) alkaline phosphatase (ALP), (C) gamma-glutamyl transferase (GGT), (D) bile acids and (E) bilirubin as well as for biomarkers of renal dysfunction such as urea (F). Each symbol represents the values for an individual sheep at the different time points. The dotted lines indicate the reference values, which were established from the maximum and minimum measurements obtained in serum samples collected before inoculation (day 0). Statistically significant differences at different days post-inoculation (mean ± SD) between subcutaneously and intranasally inoculated animals and between both groups of in-contact sheep were analysed by two-way ANOVA. Statistical significance between groups is annotated with black asterisks. Statistically significant differences within each experimental group at different days post inoculation relative to pre-inoculation values were assessed by one-way ANOVA. Statistical differences are indicated by red (intranasal group) and blue (subcutaneous group) asterisks. (x-axis): day post-inoculation (dpi); (y-axis): analytes concentrations; variables of significance (*P ≤ 0.05; **P ≤ 0.01; ***P ≤ 0.001; ****P ≤ 0.0001); SC: subcutaneous group; IN: Intranasal group.

Regarding analytes indicative of renal failure, urea levels increased moderately from 2 dpi in both groups and remained slightly elevated until the end of the experiment, especially in animals in the subcutaneous group, suggesting mild renal impairment (Fig. 6F). Creatinine concentrations did not show remarkable changes (S1 Fig).

Throughout the experiment, there were no noticeable changes in any of the inoculated groups in biomarkers indicative of metabolic disturbances, such as glucose, triglycerides and cholesterol (S1 Fig), or in other immunological and inflammatory markers, such as total protein, lactate, adenosine deaminase and haptoglobin (S2 Fig).

Among the in-contact animals, the most notable changes in hepatic biomarkers (AST, ALP, GGT, bile acids, and bilirubin) were observed in in-contact sheep #20 from the subcutaneous group (Fig. 7A-7E). This animal, together with in-contact sheep #19 from the same group, also showed mild to moderate increases in urea levels suggestive of mild renal impairment (Fig. 7F). No remarkable changes were observed in the concentrations of the other analytes assessed (S3 and S4 Fig).

**Fig. 7.**
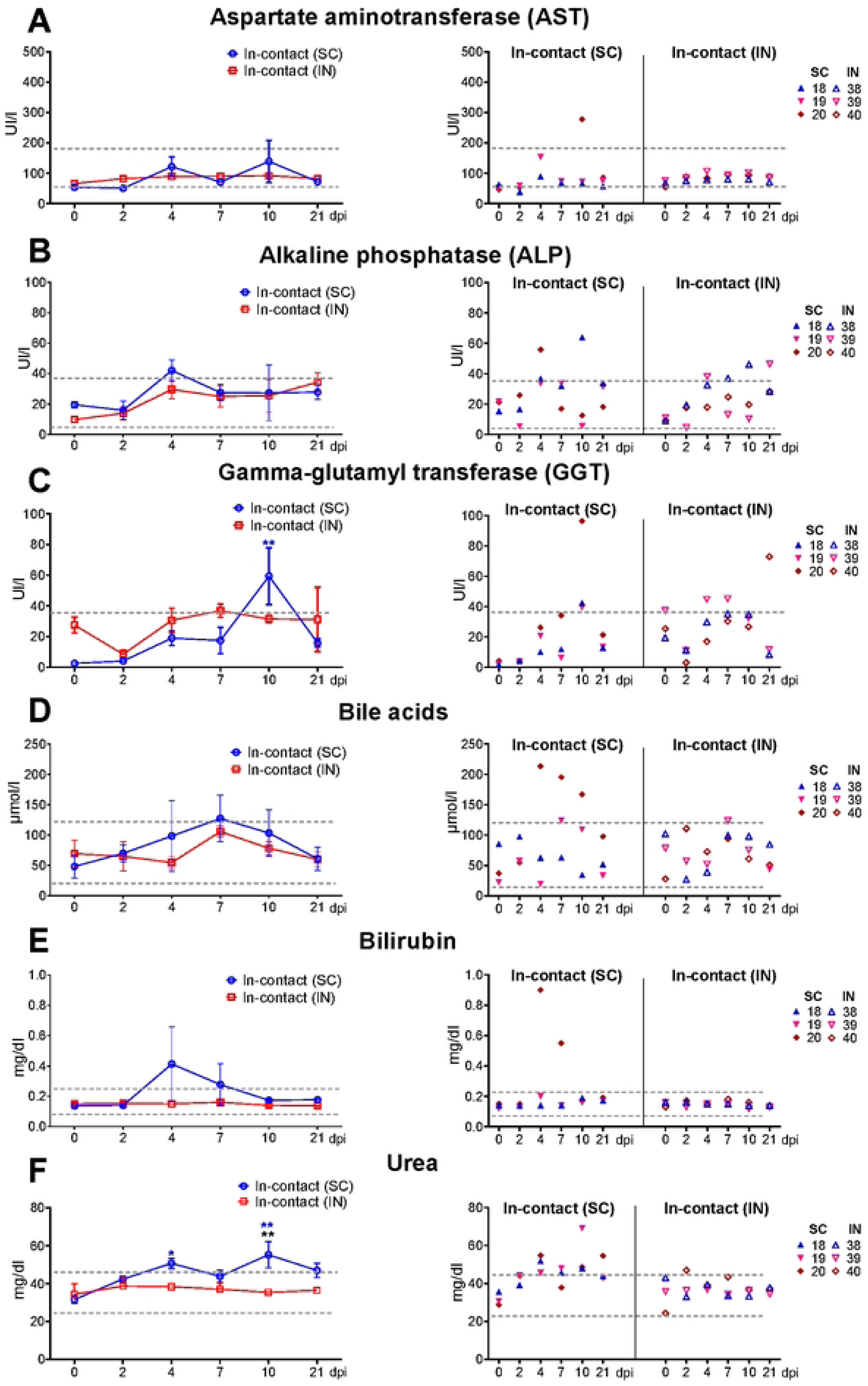
Serum clinical biochemical parameters in in-contact sheep. Serum samples from in-contact sheep housed in the subcutaneous or intranasal group were analysed for biomarkers indicative of hepatic damage, including (A) aspartate aminotransferase (AST), (B) alkaline phosphatase (ALP), (C) gamma-glutamyl transferase (GGT), (D) bile acids and (E) bilirubin as well as for biomarkers of renal dysfunction such as urea (F). Each symbol represents the values for an individual sheep at the different time points. The dotted lines indicate the reference values, which were established from the maximum and minimum measurements obtained in serum samples collected before inoculation (day 0). Statistically significant differences at different days post-inoculation (mean ± SD) between subcutaneously and intranasally inoculated animals and between both groups of in-contact sheep were analysed by two-way ANOVA. Statistical significance between groups is annotated with black asterisks. Statistically significant differences within each experimental group at different days post inoculation relative to pre-inoculation values were assessed by one-way ANOVA. Statistical differences are indicated by red (intranasal group) and blue (subcutaneous group) asterisks. (x-axis): day post-inoculation (dpi); (y-axis): analytes concentrations; variables of significance (*P ≤ 0.05; **P ≤ 0.01; ***P ≤ 0.001; ****P ≤ 0.0001); SC: subcutaneous group; IN: Intranasal group.

### Histopathological evaluation

Although no remarkable macroscopic findings were observed during necropsy, histopathological examination of the sampled organs revealed lesions in the liver and frontal cortex of the brain. In the liver, all SC and IN inoculated sheep, as well as the in-contact sheep housed in the subcutaneous group, particularly sheep #20, exhibited chronic, multifocal, mild to moderate hepatitis (S5 Fig). The lesions were characterised by mononuclear inflammatory infiltrates predominantly within the portal spaces, with occasional focal infiltrates into the hepatic parenchyma and surrounding the centrilobular vein. These infiltrates were composed predominantly of lymphocytes, with occasional plasma cells and macrophages. This histopathological pattern was indicative of persistent immune-mediated hepatic injury and aligned with the characteristic lesions of chronic viral hepatitis.

On the other hand, most inoculated sheep exhibited a non-suppurative meningoencephalitis in the frontal cortex (Fig. 8). Lesions were mild and focal in the SC inoculated sheep and consisted of low number of lymphocytic infiltrates within the meninges, accompanied by mild satellitosis and focal gliosis in the neuroparenchyma, with lymphocytes perivascular cuffing occasionally present within the Virchow–Robin spaces. In contrast, IN inoculated sheep exhibited more severe lesions, characterized by thickened meninges with higher numbers of lymphocytic infiltrates, along with diffuse satellitosis and multifocal moderate gliosis in the neuroparenchyma, as well as dense perivascular cuffs. Similar lesions, although less severe, were observed in the in-contact sheep housed in the subcutaneous group, particularly sheep #18 and #20, whereas no lesions were detected in the in-contact sheep housed in the intranasal group. These findings reflected a subacute process and displayed the typical pattern associated with viral infections of the central nervous system.

**Fig. 8.**
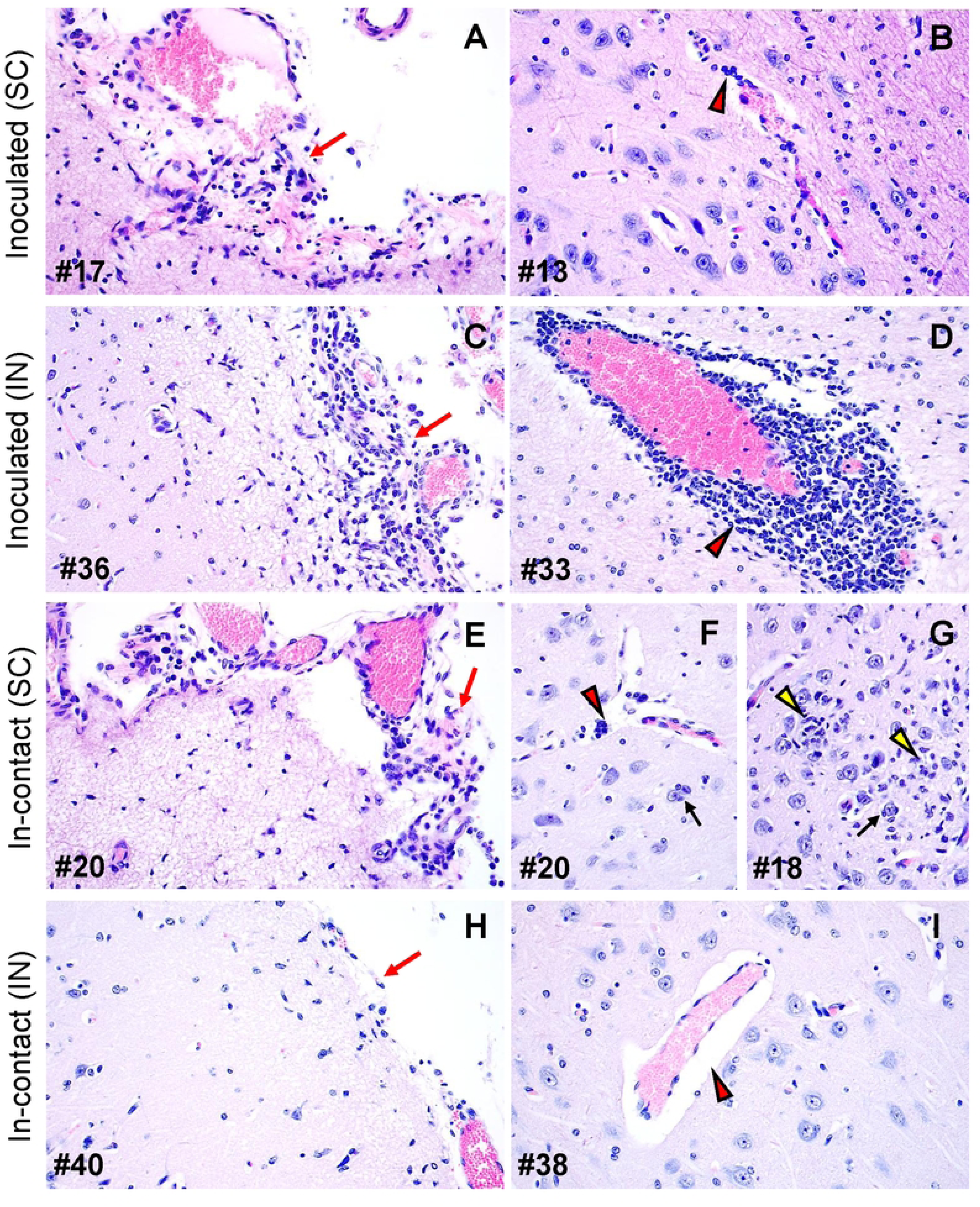
Representative histopathological images of the frontal cortex of the brain. All animals were euthanised at 28 days post-inoculation (dpi). Subcutaneously (SC) and intranasally (IN) inoculated sheep exhibited a non-suppurative meningoencephalitis in the frontal cortex. Observe the low number of lymphocytic infiltrates within the meninges **(A, C, red arrow)** which were prominent in the IN inoculated animals. Observe also the presence of focal, mild lymphocytic infiltrates within the meninges of the in-contact sheep housed in the subcutaneous group **(E, red arrow)**, in contrast to the histologically unremarkable changes in meninges of the in-contact sheep housed in the intranasal group **(H, red arrow)**. SC inoculated sheep also showed lymphocytes perivascular cuffing occasionally present within the Virchow–Robin spaces **(B, red arrowhead)**, whereas IN inoculated animals exhibited dense perivascular cuffs **(D, red arrowhead)**. In-contact sheep housed with the subcutaneous group also showed lymphocytes occasionally present within the Virchow–Robin spaces **(F, red arrowhead)**, accompanied, similar to the IN inoculated sheep, by focal gliosis **(G, yellow arrowhead)** and satellitosis **(F-G, black arrow)**. None of these findings were observed in the in-contact sheep housed in the intranasal group **(I)**. Also, most of the meningeal and neuroparenchymal capillaries and venules are frequently congested in areas with lymphocytic meningitis and intra-neuroparenchymal perivasculitis with occasional hypertrophy of the endothelial cells **(A-F)**. Haematoxylin and eosin staining; Magnification: 40×; Sheep identification number (# number).

## Discussion

The results obtained in the present study demonstrate that the route of inoculation of RVFV in sheep influences disease dynamics, antibody responses, the evolution of viremia, and the concentrations of biochemical markers indicative of liver or kidney damage. Additionally, in the absence of competent vectors, we demonstrated that the virus can be transmitted horizontally between inoculated and in-contact animals.

None of the sheep inoculated either SC or IN, the latter route of infection tested for the first time in sheep, developed clinical manifestations, apart from mild and transient increases in temperature. These results are consistent with previous experimental infections in sheep inoculated SC [27,30], as well as with studies in goats [22] and calves [23], which were inoculated IN with doses and strains of comparable virulence and likewise exhibited subclinical forms of disease. Thus, sheep, like other ruminants, showed no predisposition to developing neurological signs, in contrast to other experimental models such as mice [31–34], non-human primates [35], or ferrets [36], which exhibited viral neuroinvasion, neurological disease, and encephalitis following IN inoculation. However, the inflammatory lesions observed in the brain, described for the first time in this study, indicate central nervous system damage or dysfunction and demonstrate that the sheep inoculated IN exhibited the greatest degree of injury. However, the absence of neurological signs throughout the experiment suggests a notable resistance of sheep to clinically apparent neurological damage. Studies focusing on earlier stages of infection will be necessary to determine whether viral neuroinvasion occurred and to clarify the pathogenic mechanisms underlying the brain lesions.

Although RVFV inoculation induced only subclinical disease regardless of the route of administration, biochemical analyses revealed evidence of mild hepatic and renal damage. The dynamic of hepatic damage analytes was similar in both experimental groups, although serum concentrations of AST, ALP, and GGT were generally higher in the IN inoculated group, suggesting moderate but greater liver damage or dysfunction. This observation could be associated with the higher viremia levels observed in this group and, likely, to increased viral replication in this RVFV target organ, which was not assessed in the present study. Our findings are consistent with those of previous studies in which subcutaneous inoculation with RVFV was performed in sheep. These studies likewise reported an early rise in hepatic injury markers concentrations from 2 dpi [30,37–39]. In addition, our results also demonstrate that the inoculation route of RVFV would influence in the severity hepatic injury. Differences in the concentrations of hepatic injury markers attributable to the route of inoculation have also been suggested in cattle inoculated with RVFV by intradermal or intranasal route [23]. In terms of renal impairment, comparable trends were observed in the concentrations of urea in both groups, with modest increases from 2 dpi onwards, indicative of mild renal dysfunction. These results were consistent with those of previous subcutaneous inoculations in sheep, which also exhibited similar dynamics involving increases in blood urea nitrogen (BUN) levels from 2 dpi [30,37,38]. In cattle, only those inoculated IN with RVFV showed a mild increase in BUN concentrations from 1 dpi, whereas animals inoculated intradermally did not exhibit this change [23]. Future time-course studies using sequential euthanasia should more precisely characterise the progression, distribution and severity of RVFV-induced lesions in the aforementioned target organs across different species and inoculation routes.

Additional differences related to the inoculation route were evident in our study, with IN inoculated animals showing a delayed onset of viremia, seroconversion, and the development of nAb. However, these parameters reached higher values than those observed in SC inoculated animals. Results suggest that the intranasal route requires a longer transition period before the virus reaches the bloodstream, highlighting a potential role for mucosal immunity in delaying early viral replication, viremia, and the onset of clinical signs and systemic humoral immune responses. In addition to the higher viremia levels reached in the IN inoculated sheep, it is noteworthy that the viremia dynamics and levels observed in the in-contact sheep within the subcutaneous group closely resembled those of the IN inoculated animals. This fact may suggest that these in-contact sheep became infected via the nasal route, although other potential routes, such as the conjunctiva or the oropharynx, cannot be ruled out. Comparative studies in other ruminant species, such as goats [22] and cattle[23] have likewise reported a delayed onset of viremia, along with higher viremia titres, following intranasal inoculation compared with parenteral routes, with viremia kinetics closely resembling those observed in our study.

In cattle, a comparative study that used different inoculation routes suggested that IN inoculated animals may delay active RVFV replication at the level of the mucosa through IFNα secretion, cytokine response that was not observed in those intradermally inoculated [23]. Although early IFNα production may establish an antiviral state at the mucosal level, limiting initial viral replication and dissemination, RVFV would be able to circumvent this first line of defence by suppressing type I IFN signalling through the activity of its non-structural protein s (NSs protein), as demonstrated in *in vitro* infection studies [40,41]. This suppression of the innate immune response would facilitate further viral replication [42], allowing the virus to amplify more efficiently, mechanism that could explain the differences in viremia levels between our experimental groups.

A delayed nAb response in goats inoculated IN, compared with those inoculated SC, was also reported [22], displaying dynamics similar to those observed in the sheep involved in our experiment. In other parenteral RVFV inoculation studies in sheep, nAb were detected from 5 to 6 dpi [43,44], slightly later than in our SC inoculated group, differences that may be attributable to variations in sampling schedules and experimental design. These results demonstrate that the route of administration can shape the activation of the humoral immune response, an essential component of protection against RVFV [45–47]. This should consider when designing and administering live-attenuated RVFV vaccines [48], as the subcutaneous route appears to induce a less robust, albeit faster, neutralizing antibody response than the intranasal route. Furthermore, the intranasal route may pose greater safety concerns due to its potential to cause neurological damage as demonstrated in the present study.

Although the SC inoculated group developed lower levels of viremia compared with the IN inoculated group, only animals in contact within the SC inoculated group developed transient subclinical disease, seroconverted, and became viremic, with infectious virus also being isolated. All these changes occurred later than those observed in the SC inoculated animals housed in the same room, providing evidence of RVFV transmission and suggesting that infection via parenteral routes such as subcutaneous inoculation may create favourable conditions for RVFV secretion, thereby facilitating horizontal transmission. Previous studies in which sheep were inoculated parenterally (intravenously or subcutaneously) with RVFV strains of comparable virulence, reported dynamics and titres of viremia similar to those observed in our study. However, none of the non-inoculated sentinel or control animals in those experiments showed evidence of horizontal virus transmission [29,37,49]. In contrast, other experimental studies reported only sporadic cases of horizontal transmission to in-contact sheep from animals inoculated SC with RVFV [26–28].

In our study, infectious virus was isolated from most of the inoculated sheep between 1 and 3 dpi, regardless of the route of inoculation, with blood samples showing high viremia titres within the range (10^4.3^ to 10 ^10.2^ PFU/ml) previously suggested in theoretical prediction models as required for virus transmission by mosquito vectors [50]. In other RVFV mosquito–sheep transmission studies [51], virus was isolated only from mosquitoes fed at 2 dpi on intravenously inoculated sheep that were experiencing peak viremia levels (between 10^5.2^ to 10^5.7^ TCID₅₀/ml), comparable to those observed in our animals and coinciding with the period (1–3 dpi) during which virus isolation was successful in both experimental groups. Therefore, the presence of infectious virus coinciding with high levels of viremia in the early stages after inoculation suggests that, in the absence of competent vectors, the virus could be excreted and transmitted horizontally to uninoculated animals housed in the same room. However, such horizontal transmission was only confirmed in those in-contact animals in the subcutaneous group. Given that no infectious virus was isolated from any swabs during the experiment, and that viral genome was detected only in a few oro-nasal swabs from IN inoculated animals on days 1 and 2 post-infection, likely reflecting residual inoculum rather than active virus replication, it was not possible to determine whether rectal or oro-nasal shedding contributed to horizontal virus transmission. Therefore, the reasons why in-contact sheep in the subcutaneous group became infected, whereas those in the intranasal group did not, remain unclear. Future research should explore these mechanisms in greater depth, including the assessment of alternative excretion pathways such as ocular or urinary routes. Additionally, as highlighted by previous studies [22,52], it will be important to determine whether components present in nasal swabs, along with the transport media used may interfere with RVFV detection. Addressing these questions will help refine sampling strategies and improve diagnostic reliability.

In conclusion, SC inoculated sheep had a shorter incubation period, an earlier onset of viremia, and earlier seroconversion accompanied by an earlier neutralizing antibody response. In contrast, IN inoculated animals developed higher rectal temperatures, reached higher peak viremia, and developed a more robust neutralizing antibody response. They also exhibited increased concentrations of analytes indicative of moderate but more severe hepatic injury compared with the subcutaneous group, along with more pronounced histopathological damage in the central nervous system. However, in both experimental groups, analytes associated with renal impairment suggested only mild renal dysfunction. Our results also confirmed the horizontal transmission of RVFV between SC inoculated sheep and in-contact animals housed in the same room, pointing to the oronasal mucous membranes as the most likely route of infection. This finding not only underscores the influence of the inoculation route on virus transmission but also highlights the potentially significant role of horizontal transmission in RVF epidemiology and disease control.

## Material and methods

### Cells and virus

The virus used in our experiment was isolated in the C6/36 mosquito cell line (Aedes albopictus clone, ATCC CRL-1660™) from the plasma of a sheep infected with RVFV (strain 56/74) [53].

RVFV-56/74 stock was produced in C6/36 cells grown in Dulbecco’s modified Eagle medium (DMEM, Bio-West, Nuaillé, France) supplemented with heat inactivated 10% foetal bovine serum (FBS, Gibco™-Thermo Fisher Scientific. Waltham, MA, USA), 1% L-glutamine (2 mM), 1% non-essential amino acids, and 100 U/mL penicillin and 100 µg/mL streptomycin. Cells were incubated at 28 °C. To produce virus stock, cells were grown in T150 flasks up to 95% confluence. At this point, cells were infected with 56/74 RVFV strain with a low moi (0.01-0.05 PFU/cell) and left incubating for one hour at 28 °C. After incubation, the cells were washed twice with DMEM and fresh media was added. The cell culture was then incubated at 28 °C for three to seven days. The supernatant with virus was collected and frozen at -80 °C until titration.

For viral titration as well as for virus isolation from blood samples and swabs, Vero cell line (African green monkey kidney cells, ATCC CCL-81^TM^) was used. Cells were grown in DMEM supplemented with heat inactivated 5% foetal bovine serum, 1% L-glutamine, 1% non-essential amino-acids, and 100 U/mL penicillin and 100 µg/mL streptomycin. All cells were incubated at 37 °C in the presence of 5% CO_2_. The titration of virus stock was performed as described [54].

### Ethics statement

All animal procedures were conducted in accordance with Spanish (RD 53/2013) and European (EU Directive 2010/63/EU) regulations on the protection of animals used for scientific purposes. The experimental procedures were approved by the Animal Research Ethics Committee (Consejo Superior de Investigaciones Científicas-CSIC) and the regional authorities of the Community of Madrid (permit number PROEX/22/06.79).

### *In vivo* experimental design

Twenty Castilian breed sheep of both sexes, 9 to 10 weeks old, were obtained from a commercial farm with a high sanitary status. Following a seven-day acclimatisation period at the BSL-3 animal facility of the Centro de Investigación en Sanidad Animal (CISA-INIA/CSIC), animals were randomly assigned into two groups with ten animals each (Fig. 1). In each group, seven animals were inoculated subcutaneously (SC; numbered #11-17) or intranasally (IN; numbered #31-37) with 1 ml containing 1 × 10^6^ plaque-forming units (PFU/ml) of the virulent RVFV 56/74 strain. Intranasal mucosal atomization devices (Aluneb, MAD nasal, Cinfa) were used for intranasal inoculation. Additionally, three animals within each group (numbered #18-20 in the subcutaneous group and #38-40 in the intranasal group) were inoculated with mock-infected cell culture medium and were designated as in-contact animals.

Following inoculation, rectal temperature and clinical signs were monitored daily using clinical evaluation protocols previously described [45]. Oro-nasal and rectal swabs, as well as EDTA blood and serum samples taken from the jugular vein, were obtained from all the animals prior to infection (day 0) and at 1, 2,3,4,7, 10, 21 and 28 dpi. Blood and serum samples were stored at -80 °C. Swabs were eluted in 500 µl of sterile PBS for two hours and eluates were centrifuged at 10,000 rpm for 5 minutes at room temperature. The resulting supernatant was collected and stored at -80 °C.

At the end of the experiment (28 dpi), all animals were sedated with a combination of ketamine (Anesketin, 2 mg/kg BW) and xylazine (Xilagesic, 0.1mg/kg BW) administered intravenously. Animals were then euthanised with an overdose of intravenous pentobarbiturate (Dolethal, 20ml). All animals were necropsied, and the presence of macroscopic lesions was evaluated. Liver and spleen samples were aseptically taken and stored at -80 °C. In addition, samples from the skin at the inoculation site, pharyngeal tonsil, left cranial pulmonary lobe, liver, spleen, kidney, adrenal gland, frontal cortex of the brain, and the prescapular, tracheobronchial, and gastrohepatic lymph nodes were collected, fixed in 10% buffered formalin for 72 hours, routinely processed, and paraffin-embedded for histopathological evaluation.

### Specific and neutralising RVFV antibody detection

Antibody ELISA tests were performed on serum samples taken at 0, 3, 4, 7, 10, 21 and 28 dpi. IgM Antibody Capture (MAC) ELISA was used for the detection of anti-nucleoprotein IgM antibodies (ID Screen® Rift Valley Fever IgM Capture, Innovative Diagnostics, Grabels, France), while Competitive ELISA kit was used for the detection of anti-Rift Valley Fever antibodies (ID Screen® Rift Valley Fever Competition Multi-species, Innovative Diagnostics, Grabels, France). Protocols were performed according to manufactureŕs instructions.

The titres of RVFV serum neutralising antibodies (nAbs) were determined using a virus microneutralization test (VNT). Briefly, in a 96-well plate format, serial dilutions (50 μl) of heat-inactivated serum samples (2 hours, 56 °C), in 2–4 replicas, were incubated with 50 μl of the RVFV 56/74 strain (100 TCID_50_) for 2 hours at room temperature.

Subsequently, 2.5 × 10^4^ Vero cells (in 50 μl) were added to each well. Plates were incubated for 3–4 days at 37 °C and 5% CO_2_ and cytopathic effect (CPE) scored under microscope observation. Neutralisation titres were calculated using the Reed and Muench 50% end-point method and expressed as the reciprocal of the serum dilution that completely blocked cytopathic effect in 50% of wells (VNT_50_).

### Quantification of viral RNA levels and virus isolation in blood, swabs and tissues

RNA was extracted from EDTA blood and frozen tissue samples (liver and spleen) using the IndiSpin Pathogen Kit (Indical Bioscience), according to the manufacturers’ instructions. Prior to the RNA extraction, tissue samples were processed to obtain 10% weight/volume homogenates in sterile PBS using a Tissue lyser LT (Qiagen) by running one homogenization cycle (50 Hz oscillation frequency for 4 minutes). For swab eluates, RNA was extracted using the Speedtools RNA virus extraction kit (Biotools B&M Labs).

RT-qPCR was performed in duplicate on an Mx3000P real-time PCR system, and PFU per mL were determined by comparison to a standard curve made from RVFV 56/74 samples with a defined plaque titre. RNAemia levels were then assessed in 1 to 5 µl of extracted RNA by a RT-qPCR using the High Scriptools-Quantimix Easy Probes Kit (Biotools, Spain) and the forward RVFV L segment primer as described [55].

Attempts were made to isolate the virus *in vitro* from blood samples obtained from inoculated and in-contact animals. For each animal, the blood sample selected for virus isolation corresponded to the time point showing the highest RNAemia level. Attempts were also performed in animals in which RNAemia was not detected. In addition, virus isolation was only attempted in swabs eluates where viral genome was detected. Virus isolation was not performed in tissue samples (liver and spleen). Briefly, 100 µL of blood or swab eluates were diluted with DMEM containing 2% FBS and adsorbed for 1 hour (5% CO_2_ incubator at 37 °C) to semi confluent Vero cell monolayers in T25 flasks. After adsorption, cells were washed twice with sterile PBS and incubated with fresh medium for 5–6 days until cytopathic effect was observed; samples where cytopathic effect was not observed were subjected to two additional blind passages. Supernatant collected from CPE positive cultures were tested by indirect immunofluorescence assays to confirm the presence of RVFV in these samples.

### Serum biochemistry

Serum samples collected on days 0, 2, 4, 7, 10, and 21 dpi from sheep inoculated by both routes and from in-contact animals, were thawed and inactivated using detergent and heat (56 °C for 1 hour in the presence of 0.5% Tween 20 v/v) in accordance with WOAH guidelines [56]. Once inactivated, samples were stored at −80°C, or on dry ice during shipping, until analysed at the Laboratotio Interdisciplinar de Patología Clínica (Interlab-UMU) of the University of Murcia (Spain). Haptoglobin (Hp) concentrations were measured via a haemoglobin-binding assay using a commercially available colorimetric method (Haptoglobin Tridelta® phase range, Tridelta Development Ltd., Maynooth, Country Kildare, Ireland). The rest of the analytes were measured using an automated analyser (Olympus AU600, Olympus Diagnostic Europe GmbH, Ennis, Ireland) following the instructions of the manufacturer using Olympus commercial kits. Total serum protein, albumin, alkaline phosphatase (ALP), Gamma-glutamyl transferase (GGT), Aspartate aminotransferase (AST), bilirubin, bile acids, creatinine, urea, cholesterol, glucose, triglycerides and lactate were measured using commercial kits from Beckman Coulter; Adenosine deaminase was determined by spectrophotometry using a commercial kit from Diazyme Laboratories [57].

### Statistical analyses

All the statistical analyses and figure generation were performed using GraphPad Prism Version 8.0.1 (GraphPad Software, La Jolla, CA, USA). Statistical differences in rectal temperatures, viremia and biochemical analytes between the experimental groups at different days after inoculation (mean ± SD) were assessed using two-way ANOVA. In addition, statistical differences within each experimental group at different days after inoculation compared to pre-inoculation values were also assessed using one-way ANOVA. The statistical tests applied and variable of significance to each dataset are indicated in the corresponding figure legends.

## Acknowledgments.

We would like to thank the animal services staff, the veterinarians, the biosafety team, and the veterinary pathology staff at Centro de Investigación en Sanidad Animal (Valdeolmos, Madrid) for their valuable support with the “in vivo” experiments, and especially Antonia González Guirado, Laura Fernández del Ama and Nuria de la Losa for their excellent technical assistance. We thank Dr. Fabian Z. X. Lean (Jockey Club College of Veterinary Medicine and Life Sciences, City University of Hong Kong) for his thoughtful and critical review of the manuscript draft. We also thank to veterinary practitioner Rafael Montes-Garrido (Ganadería A.R.) for his advice and support.

## Author contributions

**Conceptualization:** Sara Morán de Bustos, Belén Borrego, Miriam Pedrera, Alejandro Brun, Belén Rodríguez-Sánchez, Pedro J. Sánchez-Cordón.

**Formal analysis:** Sara Morán de Bustos, Iris Sánchez-Del Pozo, Miriam Pedrera, José J. Cerón, Belén Borrego, Alejandro Brun, Belén Rodríguez-Sánchez, Pedro J. Sánchez-Cordón.

**Funding acquisition:** Belén Rodríguez-Sánchez, Pedro J. Sánchez-Cordón.

**Investigation:** Sara Morán de Bustos, Iris Sánchez-Del Pozo, Miriam Pedrera, José J. Cerón, Belén Borrego, Alejandro Brun, Belén Rodríguez-Sánchez, Pedro J. Sánchez-Cordón.

**Methodology:** Sara Morán de Bustos, Iris Sánchez-Del Pozo, Miriam Pedrera, José J. Cerón, Elisabeth Fuentes, David Sardón, David Rodríguez-Temporal, Belén Borrego, Alejandro Brun, Belén Rodríguez-Sánchez, Pedro J. Sánchez-Cordón.

**Project administration:** Belén Rodríguez-Sánchez, Pedro J. Sánchez-Cordón.

**Supervision:** Belén Borrego, Alejandro Brun, Belén Rodríguez-Sánchez, Pedro J. Sánchez-Cordón.

**Validation:** Sara Morán de Bustos, Iris Sánchez-Del Pozo, Belén Borrego, Alejandro Brun, Belén Rodríguez-Sánchez, Pedro J. Sánchez-Cordón.

**Visualization:** Sara Morán de Bustos, Iris Sánchez-Del Pozo, Miriam Pedrera, José J. Cerón, Belén Borrego, Alejandro Brun, Belén Rodríguez-Sánchez, Pedro J. Sánchez-Cordón.

**Writing – Original Draft Preparation:** Sara Morán de Bustos, Miriam Pedrera, José J. Cerón, Belén Borrego, Alejandro Brun, Belén Rodríguez-Sánchez, Pedro J. Sánchez-Cordón.

**Writing – Review & Editing:** Sara Morán de Bustos, Iris Sánchez-Del Pozo, Miriam Pedrera, José J. Cerón, Elisabeth Fuentes, David Sardón, David Rodríguez-Temporal, Belén Borrego, Alejandro Brun, Belén Rodríguez-Sánchez, Pedro J. Sánchez-Cordón.

## Supporting information

**S1 Fig.**
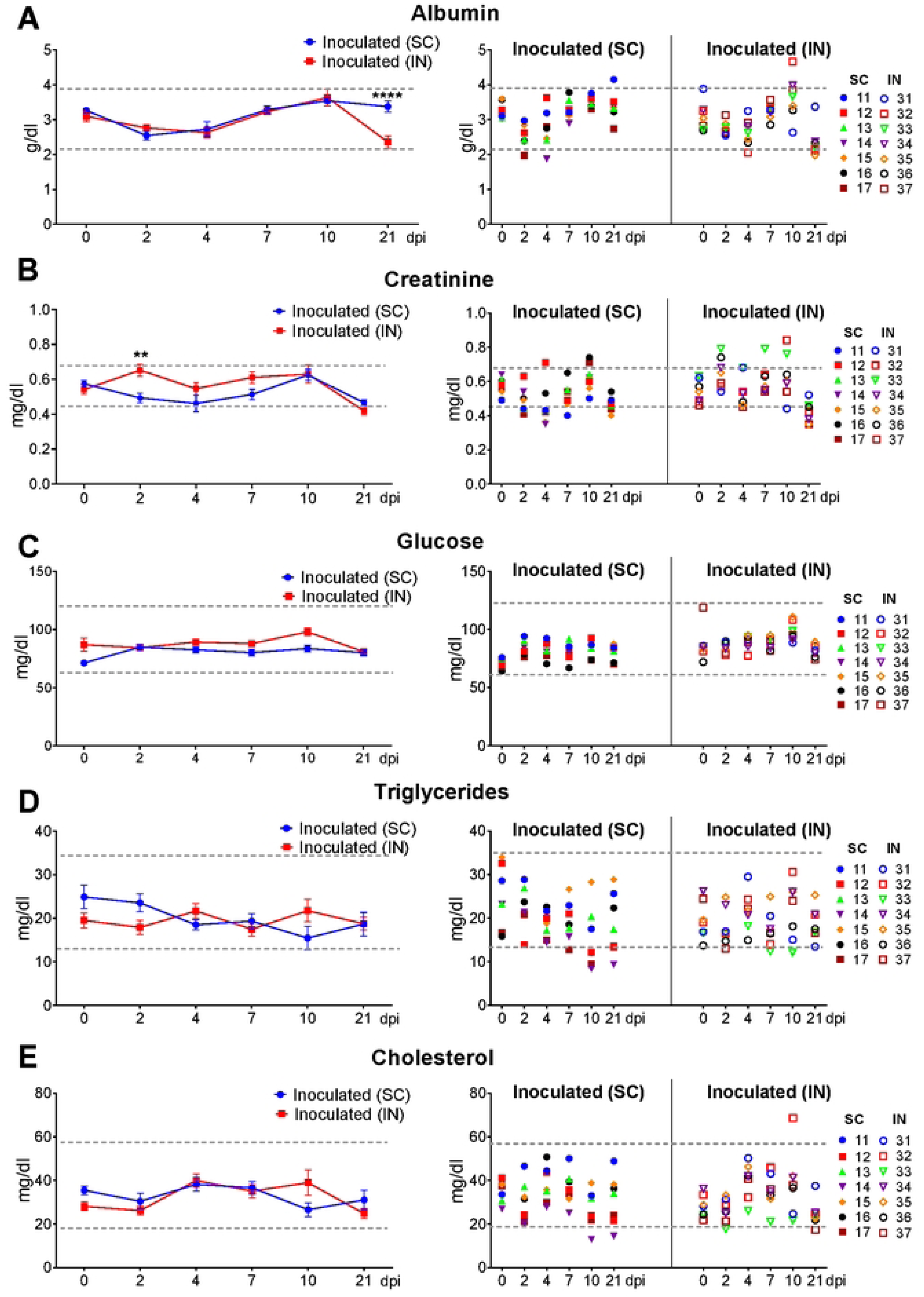
Serum clinical biochemical parameters in sheep inoculated with RVFV by subcutaneous or intranasal route. Serum samples from inoculated sheep were analysed for analytes, including (A) albumin, (B) creatinine, (C) glucose, (D) triglycerides and cholesterol (E). Each symbol represents the values for an individual sheep at the different time points. The dotted lines indicate the reference values, which were established from the maximum and minimum measurements obtained in serum samples collected before inoculation (day 0). Statistical analyses and their interpretation were conducted using the same methods described for Figure 6. (x-axis): day post-inoculation (dpi); (y-axis): analytes concentrations; SC: subcutaneous group; IN: Intranasal group.

**S2 Fig.**
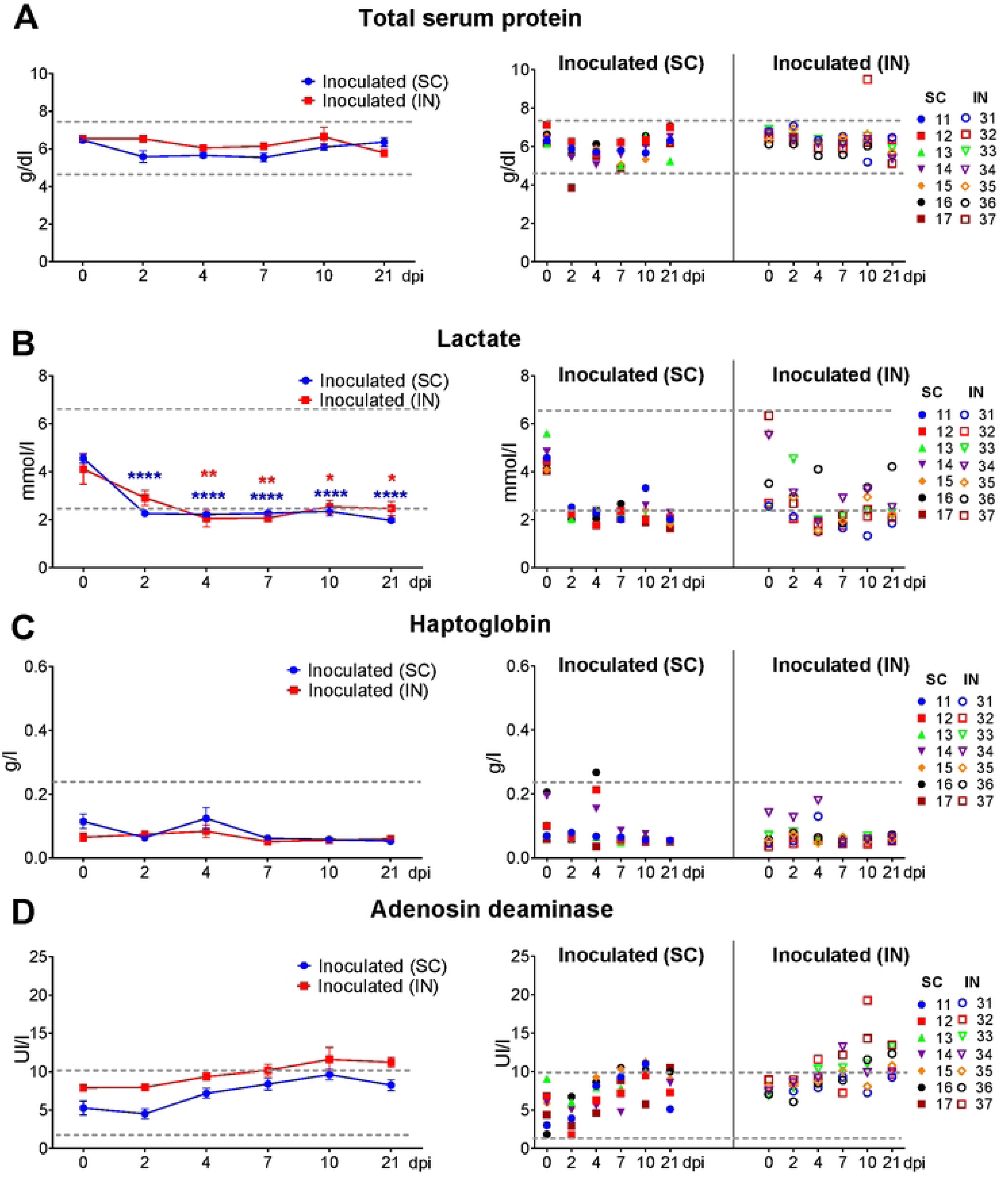
Serum clinical biochemical parameters in sheep inoculated with RVFV by subcutaneous or intranasal route. Serum samples from inoculated sheep were analysed for analytes, including (A) total serum protein, (B) lactate, (C) haptoblobin and (D) adenosine deaminase. Each symbol represents the values for an individual sheep at the different time points. The dotted lines indicate the reference values, which were established from the maximum and minimum measurements obtained in serum samples collected before inoculation (day 0). Statistical analyses and their interpretation were conducted using the same methods described for Figure 6. (x-axis): day post-inoculation (dpi); (y-axis): analytes concentrations; SC: subcutaneous group; IN: Intranasal group.

**S3 Fig.**
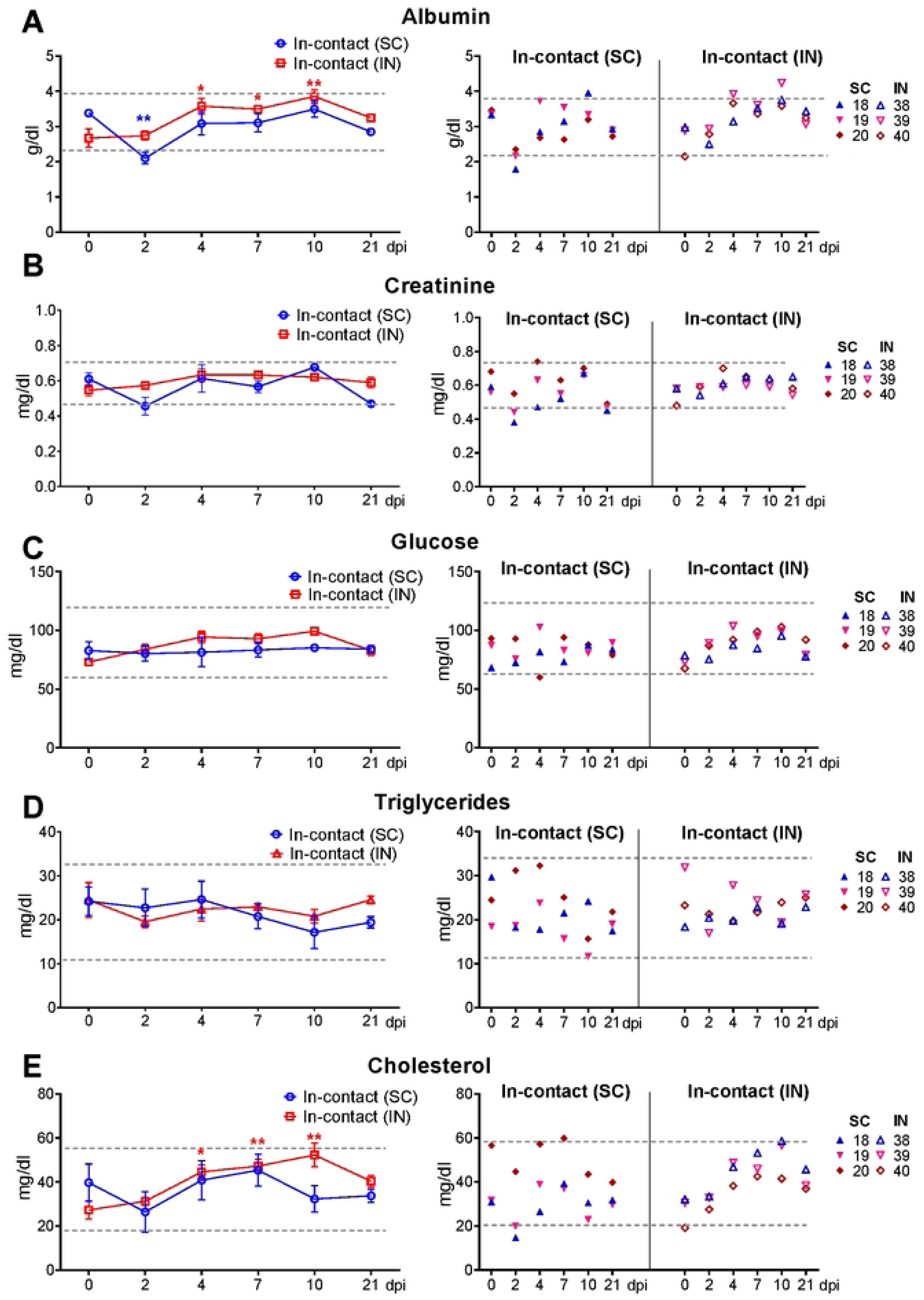
Serum clinical biochemical parameters in in-contact sheep. Serum samples from in-contact sheep were analysed for analytes, including (A) albumin, (B) creatinine, (C) glucose, (D) triglycerides and cholesterol (E). Each symbol represents the values for an individual sheep at the different time points. The dotted lines indicate the reference values, which were established from the maximum and minimum measurements obtained in serum samples collected before inoculation (day 0). Statistical analyses and their interpretation were conducted using the same methods described for Figure 6. (x-axis): day post-inoculation (dpi); (y-axis): analytes concentrations; SC: subcutaneous group; IN: Intranasal group.

**S4 Fig.**
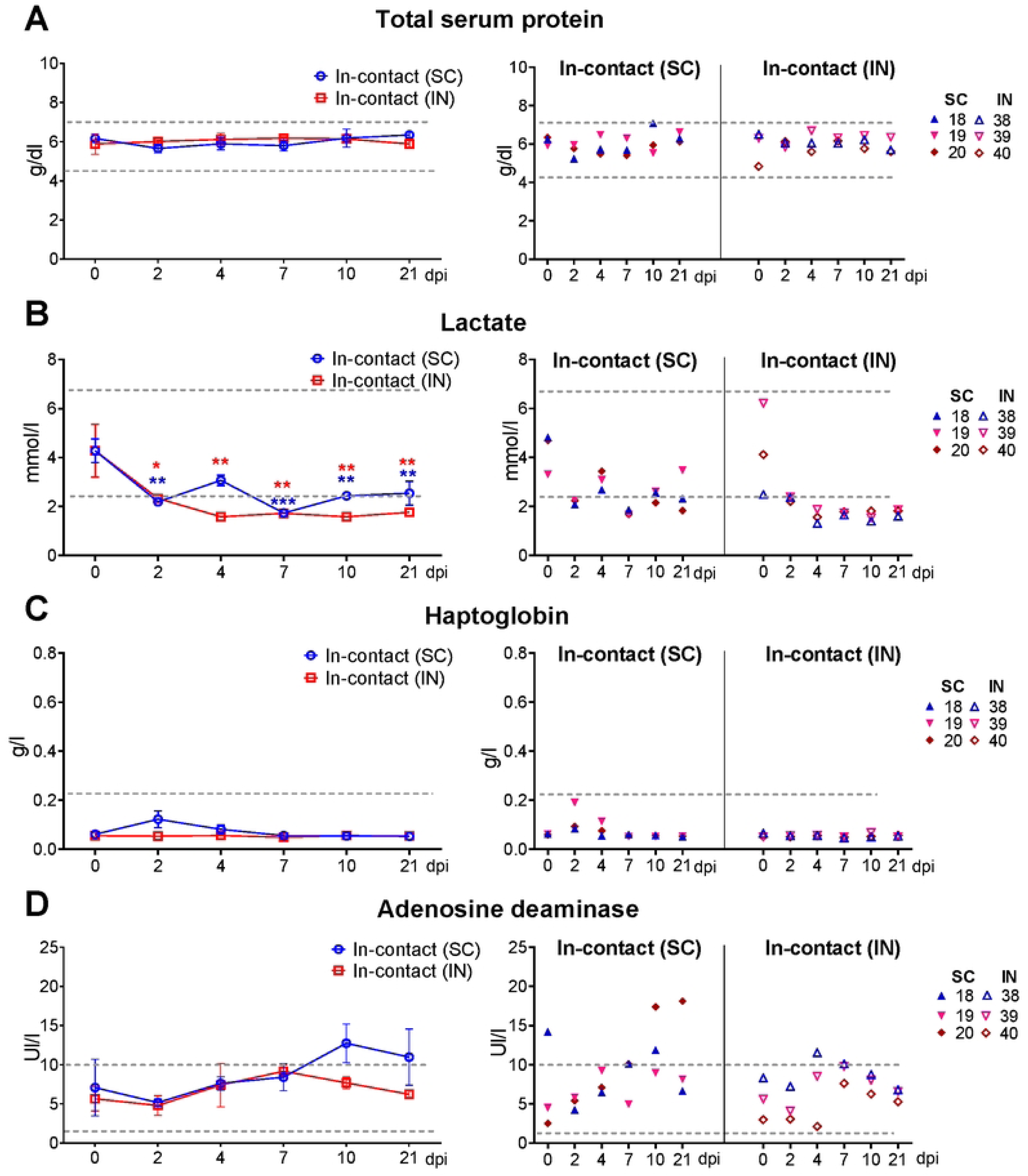
Serum clinical biochemical parameters in in-contact sheep. Serum samples from in-contact sheep were analysed for analytes, including (A) total serum protein, (B) lactate, (C) haptoblobin and (D) adenosine deaminase. Each symbol represents the values for an individual sheep at the different time points. The dotted lines indicate the reference values, which were established from the maximum and minimum measurements obtained in serum samples collected before inoculation (day 0). Statistical analyses and their interpretation were conducted using the same methods described for Figure 6. (x-axis): day post-inoculation (dpi); (y-axis): analytes concentrations; SC: subcutaneous group; IN: Intranasal group.

**S5 Fig.**
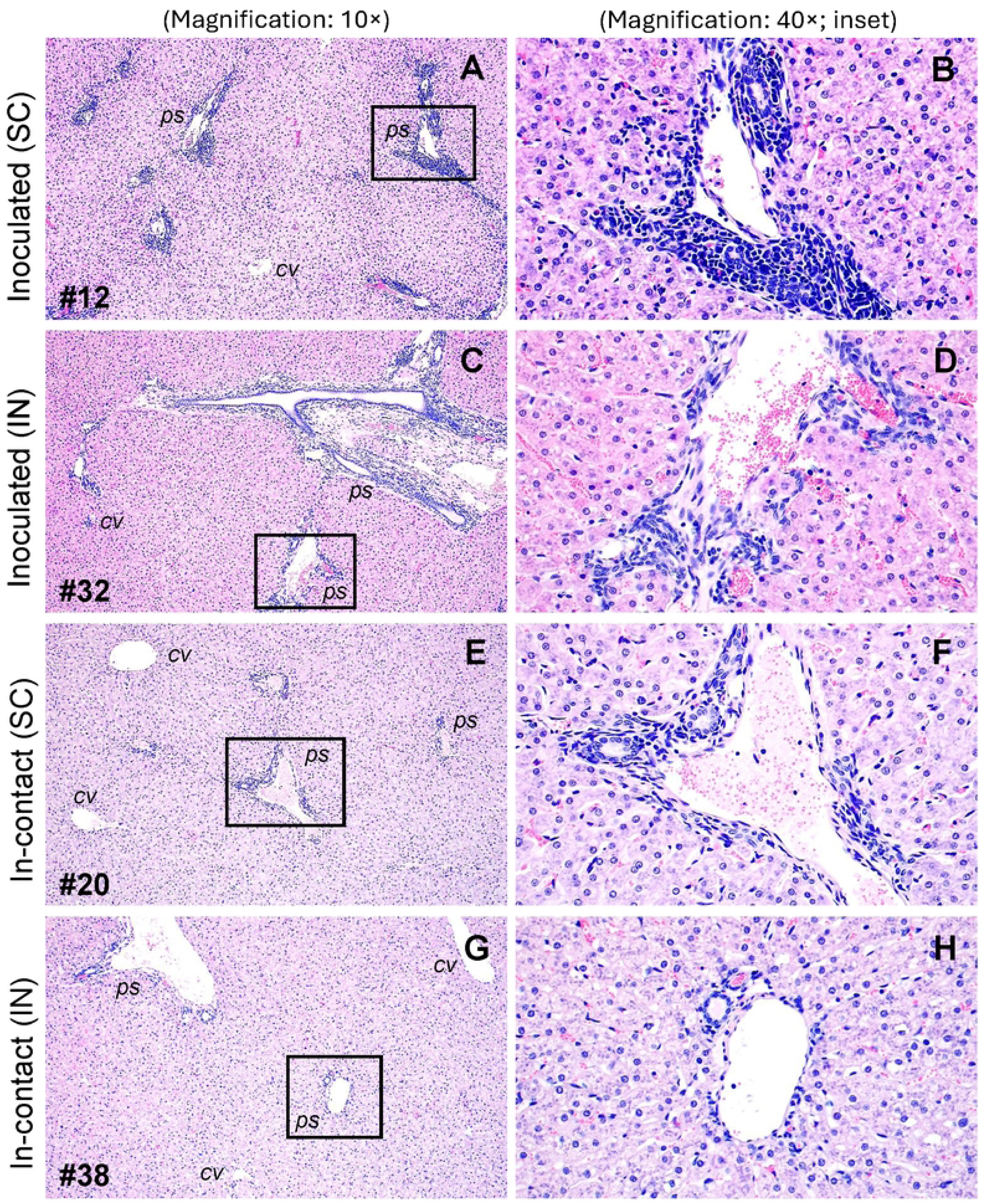
Representative histopathological images of the liver. All animals were euthanised at 28 days post inoculation (dpi). A general overview of the liver sections (magnification: 10×) is shown, accompanied by higher-magnification images of selected areas (magnification: 40×) highlighting the portal spaces (indicated by black boxes). Subcutaneously (SC) and intranasally (IN) inoculated sheep **(A, C),** as well as the in-contact sheep housed in the subcutaneous group **(E)**, exhibited chronic, multifocal, mild to moderate hepatitis characterised by mononuclear inflammatory infiltrates predominantly within the portal spaces. These infiltrates were composed predominantly of lymphocytes, with occasional plasma cells and macrophages **(B, D, F)**. These findings were occasional and mild in the in-contact sheep housed in the intranasal group **(G-H).** Haematoxylin-eosin staining; (*cv*) centrilobular vein; (*ps*) portal space; Sheep identification number (# number).

**S1 Table.** RVFV isolation from blood and oro-nasal swabs. Attempts were made to isolate the virus *in vitro* from blood samples obtained from inoculated and in-contact animals. For each animal, the blood sample selected for virus isolation corresponded to the time point showing the highest RNAemia level. Attempts were also performed in animals in which RNAemia was not detected. In addition, virus isolation was only attempted in swabs eluates where viral genome was detected. Day post-inoculation (dpi); indirect immunofluorescence (IFI).

| Sheep | Inoculated/<br>In-contact | Dpi selected for virus<br>isolation | Ct values | Cytopathic effect | IFI |
| --- | --- | --- | --- | --- | --- |
| Blood samples from sheep in the subcutaneous group |  |  |  |  |  |
| 11 | Inoculated | 1 | 34,315 | No | Negative |
| 12 | Inoculated | 2 | 27 | Yes | Positive |
| 13 | Inoculated | 1 | 37,84 | No | Negative |
| 14 | Inoculated | 2 | 27,505 | Yes | Positive |
| 15 | Inoculated | 2 | 29,74 | Yes | Positive |
| 16 | Inoculated | 1 | 34,87 | Yes | Positive |
| 17 | Inoculated | 1 | 40 | No | Negative |
| 18 | In-contact | 3 | 30,79 | Yes | Positive |
| 19 | In-contact | 3 | 33,19 | No | Negative |
| 20 | In-contact | 4 | 25,225 | Yes | Positive |
| Blood samples from sheep in the intranasal group |  |  |  |  |  |
| 31 | Inoculated | 3 | 24,93 | Yes | Positive |
| 32 | Inoculated | 3 | 28,95 | Yes | Positive |
| 33 | Inoculated | 2 | 32,335 | No | Negative |
| 34 | Inoculated | 2 | 30,745 | Yes | Positive |
| 35 | Inoculated | 2 | 26,28 | Yes | Positive |
| 36 | Inoculated | 2 | 26,955 | Yes | Positive |
| 37 | Inoculated | 2 | 28,685 | Yes | Positive |
| 38 | In-contact | 2 | 40 | No | Negative |
| 39 | In-contact | 2 | 40 | No | Negative |
| 40 | In-contact | 2 | 40 | No | Negative |
| Oro-nasal swabs from sheep in the intranasal group |  |  |  |  |  |
| 33 | Inoculated | 1 | 34,52 | No | Negative |
| 35 | Inoculated | 1 | 31,475 | No | Negative |
| 36 | Inoculated | 1 | 33,815 | No | Negative |
| 37 | Inoculated | 1 | 36,385 | No | Negative |

